# Natural depletion of H1 in sex cells causes DNA demethylation, heterochromatin decondensation and transposon activation

**DOI:** 10.1101/451930

**Authors:** Shengbo He, Martin Vickers, Jingyi Zhang, Xiaoqi Feng

## Abstract

Transposable elements (TEs), the movement of which can damage the genome, are epigenetically silenced in eukaryotes. Intriguingly, TEs are activated in the sperm companion cell – vegetative cell (VC) – of the flowering plant *Arabidopsis thaliana*. However, the extent and mechanism of this activation are unknown. Here we show that about 100 heterochromatic TEs are activated in VCs, mostly by DEMETER-catalyzed DNA demethylation. We further demonstrate that DEMETER access to some of these TEs is permitted by the natural depletion of linker histone H1 in VCs. Ectopically expressed H1 suppresses TEs in VCs by reducing DNA demethylation and via a methylation-independent mechanism. We demonstrate that H1 is required for heterochromatin condensation in plant cells and show that H1 overexpression creates heterochromatic foci in the VC progenitor cell. Taken together, our results demonstrate that the natural depletion of H1 during male gametogenesis facilitates DEMETER-directed DNA demethylation, heterochromatin relaxation, and TE activation.

## Introduction

Large proportions of most eukaryotic genomes are comprised of transposable elements (TEs), mobile genetic fragments that can jump from one location to another. For example, TEs comprise approximately 50% of the human genome (Lander et al., 2001; Venter et al., 2001), and more than 85% of the genomes in crops such as wheat and maize (Schnable et al., 2009; Wicker et al., 2018). Regarded as selfish and parasitic, activities of TEs compromise genome stability, disrupt functional genes, and are often associated with severe diseases including cancers in animals (Anwar, Wulaningsih, & Lehmann, 2017). To safeguard genome integrity, eukaryotic hosts have evolved efficient epigenetic mechanisms, including DNA methylation, to suppress TEs (He, Chen, & Zhu, 2011; Law & Jacobsen, 2010). Curiously, recent studies point to episodes of TE activation that occur in specific cell types and/or particular developmental stages (Garcia-Perez, Widmann, & Adams, 2016; Martinez & Slotkin, 2012). These TE activation events provide unique opportunities to understand epigenetic silencing mechanisms, and the co-evolution between TEs and their hosts.

Developmental TE activation has been shown in mammalian embryos, germlines and brain cells. In pre-implantation embryos and the fetal germline, LINE-1 retrotransposons are highly expressed despite relatively low levels of transposition (Fadloun et al., 2013; Kano et al., 2009; Percharde, Wong, & Ramalho-Santos, 2017; Richardson et al., 2017). Recently, LINE-1 RNA was shown to play a key regulatory role in promoting pre-implantation embryo development in mice (Percharde et al., 2018). LINE-1 elements have also been shown to transcribe and mobilise in neuronal precursor cells in mice and human (Coufal et al., 2009; Muotri et al., 2005). The underlying mechanism of such cell-specific TE activation is still unclear. Hypomethylation at LINE-1 promoters in neurons has been proposed to contribute (Coufal et al., 2009), and possibly the availability of transcription factors (Muotri et al., 2005; Richardson, Morell, & Faulkner, 2014). The frequency of LINE-1 retrotransposition in mammalian brain is still under debate, however, it is conceivable that LINE-1 activities may serve to promote genetic diversity among cells of a highly complex organ like the brain (Garcia-Perez et al., 2016; Richardson et al., 2014; Singer, McConnell, Marchetto, Coufal, & Gage, 2010).

One of the best demonstrated cases of developmental TE activation occurs in the male gametophyte of flowering plants, pollen grains. Pollen are products of male gametogenesis, which initiates from haploid meiotic products called microspores. Each microspore undergoes an asymmetric mitosis to generate a bicellular pollen comprised of a large vegetative cell (VC) and a small generative cell engulfed by the VC (Berger & Twell, 2011). Subsequently the generative cell divides again mitotically to produce two sperm. Upon pollination, the VC develops into a pollen tube to deliver the sperm to meet the female cells, and subsequently degenerates. In the mature tricellular pollen of *Arabidopsis thaliana*, several TEs were found activated and transposed (Slotkin et al., 2009). Enhancer/gene trap insertions into TEs showed specific reporter activity in the VC, and TE transpositions detected in pollen were absent in progeny (Slotkin et al., 2009). These results demonstrated TE activation in pollen is confined to the VC. TE expression in the short-lived VC has been proposed to promote the production of small RNAs, which may be transported into sperm to reinforce the silencing of cognate TEs (Calarco et al., 2012; Ibarra et al., 2012; Martinez, Panda, Kohler, & Slotkin, 2016; Slotkin et al., 2009). However, TE transcription in the VC has not been comprehensively investigated, hindering our understanding of this phenomenon.

The mechanisms underlying TE reactivation in the VC are also unknown. One proposed mechanism is the absence of the Snf2 family nucleosome remodeler DDM1 (Slotkin et al., 2009). DDM1 functions to overcome the impediment of nucleosomes and linker histone H1 to DNA methyltransferases (Lyons & Zilberman, 2017; Zemach et al., 2013). Loss of DDM1 leads to DNA hypomethylation and massive TE derepression in somatic tissues (Jeddeloh, Stokes, & Richards, 1999; Lippman et al., 2004; Tsukahara et al., 2009; Zemach et al., 2013). However, global DNA methylation in the VC is comparable to that of microspores and substantially higher than in somatic tissues (Calarco et al., 2012; Hsieh et al., 2016; Ibarra et al., 2012). This suggests that DDM1 is present during the first pollen mitosis that produces the VC, so its later absence is unlikely to cause TE activation.

A plausible mechanism underlying TE activation in the VC is active DNA demethylation. DNA methylation in plants occurs on cytosines in three sequence contexts: CG, CHG and CHH (H=A, C or T). Approximately ten thousand loci – predominantly TEs – are hypomethylated in the VC, primarily in the CG context and to a lesser extent in the CHG/H contexts (Calarco et al., 2012; Ibarra et al., 2012). Hypomethylation in the VC is caused by a DNA glycosylase called DEMETER (DME) (Ibarra et al., 2012). DME demethylates DNA via direct excision of methylated cytosine, and its expression is strictly confined to the VC and its female counterpart, the central cell, during sexual reproduction (Choi et al., 2002; Schoft et al., 2011). DME demethylation may therefore cause TE transcription in the VC, however, this hypothesis has not been tested.

Another plausible mechanism for epigenetic TE activation is chromatin decondensation (Feng, Zilberman, & Dickinson, 2013). Drastic reprogramming of histone variants and histone modifications occurs during both male and female gametogenesis, rendering the gametes and companion cells with radically different chromatin states (Baroux, Raissig, & Grossniklaus, 2011; Borg & Berger, 2015). For example, centromeric repeats, which are condensed in sperm and other cell types, are decondensed in the VC, accompanied by the depletion of centromeric histone H3 (Ingouff et al., 2010; Merai et al., 2014; Schoft et al., 2009). Chromocenters, which are comprised of condensed pericentromeric heterochromatin and rDNA repeats (Chandrasekhara, Mohannath, Blevins, Pontvianne, & Pikaard, 2016; Fransz, de Jong, Lysak, Castiglione, & Schubert, 2002; Tessadori, van Driel, & Fransz, 2004), are observed in sperm nuclei but absent in the VC nucleus, suggesting that pericentromeric heterochromatin is decondensed in the VC (Baroux et al., 2011; Ingouff et al., 2010; Schoft et al., 2009). Heterochromatin decondensation in the VC is proposed to promote rDNA transcription that empowers pollen tube growth (Merai et al., 2014). However, the cause of VC heterochromatin decondensation remains unclear.

Our previous work showed that histone H1, which binds to the nucleosome surface and the linker DNA between two adjacent nucleosomes (Fyodorov, Zhou, Skoultchi, & Bai, 2018), is depleted in *Arabidopsis* VC nuclei (Hsieh et al., 2016). H1 depletion in the VC has also been observed in a distantly related lily species (Tanaka, Ono, & Fukuda, 1998), suggesting a conserved phenomenon in flowering plants. In *Drosophila* and mouse embryonic stem cells, H1 has been shown to contribute to heterochromatin condensation (Cao et al., 2013; Lu et al., 2009). H1 is also more abundant in heterochromatin than euchromatin in *Arabidopsis* (Ascenzi & Gantt, 1999; Rutowicz et al., 2015). However, it is unknown whether H1 participates in heterochromatin condensation in plant cells, and specifically whether the lack of H1 contributes to heterochromatin decondensation in the VC.

Whether and how the depletion of H1 in the VC contributes to TE derepression is also unclear. A recent study pointed to an intriguing link between H1 and DME. In the central cell, the histone chaperone FACT (facilitates chromatin transactions) is required for DME-directed DNA demethylation in heterochromatic TEs, and this requirement is dependent on H1 (Frost et al., 2018). However, DME activity in the VC is independent of FACT (Frost et al., 2018). One attractive hypothesis is that the lack of H1 in the VC causes heterochromatin decondensation and thereby contributes to the independence of DME from FACT. H1 depletion may therefore participate in VC TE activation by promoting DME-directed demethylation. Additionally, H1 depletion may activate TE transcription independently of DNA methylation, as shown in *Drosophila* where DNA methylation is absent (Iwasaki et al., 2016; Lu et al., 2013; Vujatovic et al., 2012; Zemach, McDaniel, Silva, & Zilberman, 2010; G. Q. Zhang et al., 2015).

In this study, we identify heterochromatic TEs that are epigenetically activated in *Arabidopsis* VCs. We demonstrate that these TEs are typically subject to DME-directed demethylation at the transcriptional start site (TSS), which is at least partially permitted by the depletion of H1. However, we find that loss of H1 activates some TEs without altering DNA methylation. We also show that developmental depletion of H1 decondenses heterochromatin in late microspores and is important for pollen fertility. Our results demonstrate that H1 condenses heterochromatin in plants and maintains genome stability by silencing TEs via methylation-dependent and ‐independent mechanisms.

## Results

### Heterochromatic transposons are preferentially expressed in the vegetative cell

To measure the extent of TE activation in the VC, we performed RNA-seq using mature pollen grains, followed by the annotation of gene and TE transcripts using Mikado and the TAIR10 annotation (Venturini, Caim, Kaithakottil, Mapleson, & Swarbreck, 2018). We identified 114 TEs that are transcribed at significantly higher levels in pollen than rosette leaves (fold change > 2; p<0.05, likelihood ratio test), and hence likely to be specifically activated in the VC (*Figure 1—source data 1*) (Slotkin et al., 2009).

The VC-activated TEs are primarily located in pericentromeric regions and exhibit features of heterochromatic TEs, such as being long and GC rich (Frost et al., 2018) (*Figure 1A,B*, *Figure 1—figure supplement 1A*). As is typical of heterochromatic TEs (Zemach et al., 2013), VC-activated TEs are significantly enriched in dimethylation of histone H3 on lysine 9 (H3K9me2) in somatic tissues, and are significantly depleted of euchromatin-associated modifications (*Figure 1B, Figure 1—figure supplement 1B*). VC-activated TEs encompass diverse TE families, among which MuDR DNA transposons and Gypsy LTR-retrotransposons are significantly overrepresented (p<10^−9^ and 0.01, respectively, Fisher’s exact test; *Figure 1C*).

**Figure 1.**
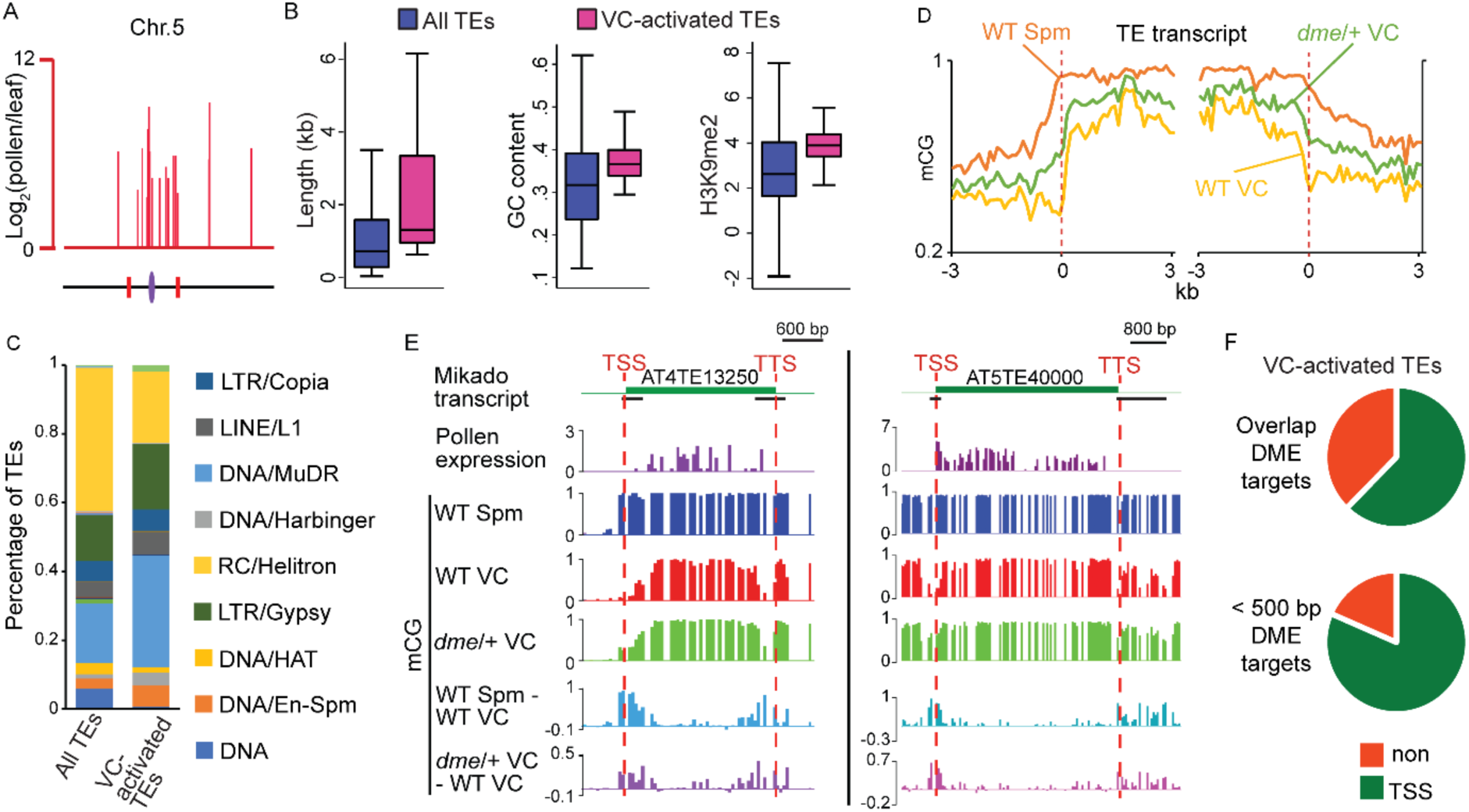
VC-activated TEs are heterochromatic and demethylated by DME. (**A**) Expression and locations of VC-activated TEs along Chromosome 5. The purple ellipse and red bars indicate the centromere and borders of pericentromeric regions, respectively. (**B**) Box plots showing the length, GC content, and H3K9me2 level of TEs. Each box encloses the middle 50% of the distribution, with the horizontal line marking the median and vertical lines marking the minimum and maximum values that fall within 1.5 times the height of the box. Difference between the two datasets compared for each feature is significant (Kolmogorov-Smirnov test p<0.001). (**C**) Percentages of TEs classified by superfamily. (**D**) VC-activated TEs were aligned at the TSS and TTS (dashed lines), respectively, and average CG methylation levels for each 100-bp interval were plotted (referred to as ends analysis). (**E**) Snapshots demonstrating the expression (Log_2_RPKM), absolute and differential CG methylation at two example VC-activated TEs. Black lines under TE annotations indicate VC DME targets. (**F**) Pie charts illustrating percentages of VC-activated TEs with TSS overlapping (top) or within 500 bp (bottom) of VC DME targets. Spm, sperm. The following figure source data and supplement are available for Figure 1: **Figure 1—source data 1.** List of VC-activated TEs. **Figure 1—source data 2.** List of VC DME targets. **Figure 1—figure supplement 1.** VC-activated TEs are heterochromatic and demethylated by DME.

### Transposon derepression in the VC is caused by DME-directed DNA demethylation

To assess whether TE activation in the VC is caused by DME-mediated DNA demethylation, we examined DNA methylation in VC and sperm at the 114 activated TEs. We found that these TEs have substantially lower CG methylation in the VC than in sperm at and near the TSS (*Figure 1D,E*, *Figure 1—figure supplement 1D*), indicative of DME activity. Because TEs tend to be flanked by repeats (Joly-Lopez & Bureau, 2018), the transcriptional termination site (TTS) regions of activated TEs also tend to be hypomethylated in the VC (*Figure 1D,E*, *Figure 1— figure supplement 1D*). Examination of DNA methylation in VCs from *dme/+* heterozygous plants (*dme* homozygous mutants are embryonic lethal), which produce a 50:50 ratio of *dme* mutant and WT pollen, revealed partial restoration of methylation at TSS and TTS of VC-activated TEs (*Figure 1D,E)*. CHG and CHH methylation is also substantially increased at the TSS (and TTS) of VC-activated TEs in *dme*/+ VC (*Figure 1—figure supplement 1C*), consistent with the knowledge that DME demethylates all sequence contexts (Gehring et al., 2006; Ibarra et al., 2012).

Consistent with the above results, 71 of the 114 (62%) VC-activated TEs overlap VC DME targets at their TSSs (*Figure 1F, Figure 1—source data 1 &2*). 96 out of the 114 TEs (84%) have VC DME targets within 500 bp of the TSS (*Figure 1F, Figure 1—source data 1*). As DNA methylation at/near the TSS has been well-demonstrated to suppress the transcription of genes and TEs in plants and animals (Barau et al., 2016; Eichten et al., 2012; Hollister & Gaut, 2009; Manakov et al., 2015; Meng et al., 2016), our results indicate that DME-directed demethylation is a major mechanism of TE activation in the VC.

### Vegetative-cell-expressed H1 impedes DME from accessing heterochromatic transposons

We next tested our hypothesis that the lack of histone H1 in the VC (Hsieh et al., 2016) allows heterochromatin to be accessible by DME. We first examined the developmental timing of H1 depletion during microspore and pollen development using GFP translational fusion lines (Hsieh et al., 2016; She et al., 2013). There are three H1 homologs in *Arabidopsis*, with H1.1 and H1.2 encoding the canonical H1 proteins, and H1.3 expressed at a much lower level and induced by stress (Rutowicz et al., 2015). H1.1- and H1.2- GFP reporters exhibit the same expression pattern: present in early microspore nucleus but absent in the late microspore stage, and remaining absent in the VC nucleus while present in the generative cell and subsequent sperm nuclei (*Figure 2A*). H1.3 is not detectable in either microspore or pollen (*Figure 2A*). These results are consistent with our previous observations, confirming that H1 is absent in the VC (Hsieh et al., 2016), and demonstrating that H1 depletion begins at the late microspore stage.

**Figure 2.**
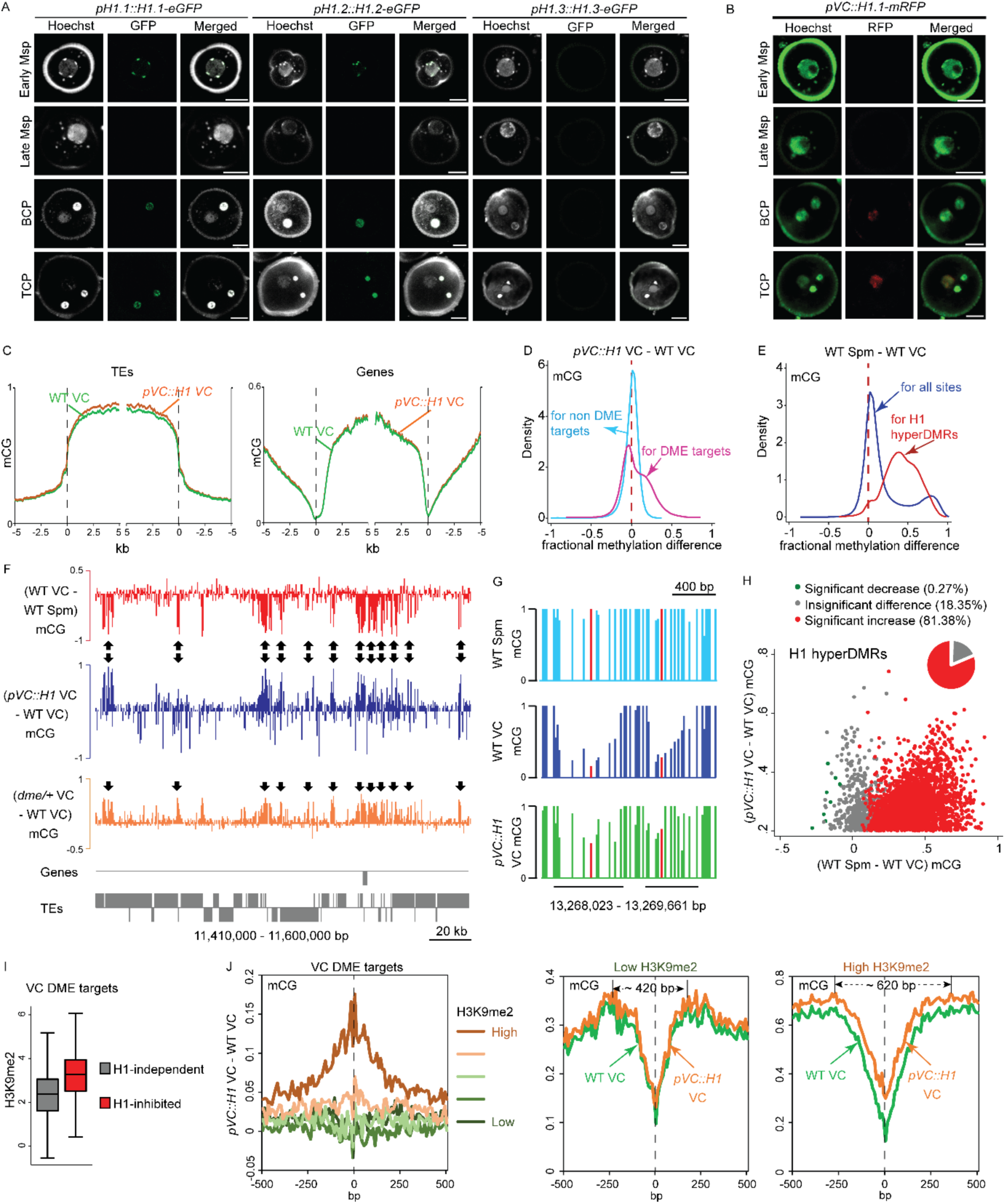
Ectopic H1 expression in the vegetative cell impedes DME at the most heterochromatic loci. (**A–B**) Confocal images showing H1 localization under native promoter (**A**) and VC-specific promoter (*pVC,* **B**) during male gametogenesis. Msp, microspore; BCP, bicellular pollen; TCP, tricellular pollen. Bars, 5 μm. All *pVC::H1.1-mRFP* (short as *pVC::H1*) refers to line #2. (**C**) Ends analysis of all TEs or genes in VCs from *pVC::H1* (line #2) and WT. (**D–E**) Kernel density plots illustrating frequency distribution of methylation differences in 50bp windows between VCs from *pVC::H1* and WT (**D**), and between WT sperm (Spm) and VC (**E**). (**F**) Snapshots showing CG methylation difference between the indicated cell types. Arrows point to DME targets that are hypermethylated by *pVC::H1*. (**G**) Snapshots demonstrating CG methylation in sperm and VCs at single-nucleotide resolution, with the cytosine most hypomethylated by DME marked in red. VC DME targets are underlined in black. (**H**) Scatter plot illustrating CG methylation differences between the indicated cell types at H1 hyperDMRs. 82.25% of H1 hyperDMRs show significant increase in sperm in comparison to VCs. (**I**) Box plot illustrating H3K9me2 level at VC DME targets that are significantly hypermethylated in *pVC::H1* (H1-inhibited) or not (H1-independent), respectively. Difference between the two groups is significant (Kolmogorov-Smirnov test p < 0.001). (**J**) VC DME targets were grouped according to H3K9me2 levels, aligned at the most demethylated cytosine (dashed lines), and plotted for average CG methylation difference as indicated in each 10-bp interval (left). Similarly, CG methylation in *pVC::H1* and WT VCs was plotted for the group with the lowest and highest H3K9me2, respectively. Spm, sperm. The following figure source data and supplements are available for Figure 2: **Figure 2—source data 1.** List of H1 hyperDMRs. **Figure 2—figure supplement 1.** H1 ectopic expression in the vegetative cell causes pollen defect and reduced fertility. **Figure 2—figure supplement 2.** H1 ectopic expression in the vegetative cell causes DNA hypermethylation at DME targets.

To understand how H1 affects DME activity, we ectopically expressed H1 in the VC. To ensure H1 incorporation into VC chromatin, we used the *pLAT52* promoter, which is expressed from the late microspore stage immediately prior to Pollen Mitosis 1, and is progressively upregulated in VC during later stages of pollen development (Eady, Lindsey, & Twell, 1994; Grant-Downton et al., 2013). Using *pLAT52* to drive the expression of H1.1 tagged with mRFP (simplified as *pVC::H1*), we observed continuous H1-mRFP signal in the VC at the bicellular and tricellular pollen stages, while the signal was undetectable in the generative cell and sperm (*Figure 2B*). H1-mRFP signal was also undetectable in late microspores (*Figure 2B*), probably due to the low activity of *pLAT52* at this stage (Eady et al., 1994). Notably, we found H1 expression in VC leads to shortened siliques and a substantial proportion of malformed pollen (*Figure 2—figure supplement 1A-C*), suggesting the depletion of H1 in the VC is important for pollen fertility.

To evaluate the effect of VC-expressed H1 on DNA methylation, we obtained genome-wide methylation profiles for VC nuclei from a strong *pVC::H1* line (#2; *Figure 2B*) and WT via fluorescence-activated cell sorting (FACS) followed by bisulfite sequencing (Supplementary file 1). CG methylation in the VC of *pVC::H1* plants is largely similar to that of WT, except for a slight increase in TE methylation (*Figure 2C*, *Figure 2—figure supplement 2A*). Consistently, the frequency distribution of CG methylation differences between VCs of *pVC::H1* and WT at loci that are not DME targets peaks near zero, showing almost no global difference (*Figure 2D*). However, a substantial proportion of loci that are targeted by DME show hypermethylation in *pVC::H1* VC (*Figure 2D*). DME targets also show preferential hypermethylation in CHG and CHH contexts in the VC of *pVC::H1* (*Figure 2—figure supplement 2B-C*). These results indicate that H1 expression in the VC specifically impedes DME activity.

**Supplementary file 1.** Sequencing summary statistics for bisulfite sequencing libraries. Mean DNA methylation (Met) was calculated by averaging methylation of individual cytosines in each context, and chloroplast CHH methylation was used as a measure of cytosine non-conversion and other errors. SN, sperm nuclei; VN, vegetative nuclei.

| Sample | Nuclear Genome Coverage | Nuclear CG Met | Nuclear CHG Met | Nuclear CHH Met | Chloroplast CHH Met |
| --- | --- | --- | --- | --- | --- |
| WT SN | 37.0 | 24.7% | 8.0% | 1.1% | 0.33% |
| WT VN | 43.7 | 14.4% | 5.2% | 1.8% | 0.45% |
| *pVC::H1* VN | 62.6 | 16.1% | 5.8% | 1.9% | 0.53% |

Across the genome, we found 2964 differentially methylated regions (DMRs) that are significantly CG hypermethylated in the VC of *pVC::H1* plants (referred to as H1 hyperDMRs hereafter; ranging from 101 to 2155 nt in length, 280 nt on average; *Figure 2—source data 1*). Most of the H1 hyperDMRs (1618, 55%) overlap DME targets in the VC (*Figure 2—source data 1*), and H1 hyperDMRs exhibit strong hypomethylation in WT VCs, with 81.4% (2412 sites) having significantly more CG methylation in sperm than VC (p<0.001, Fisher’s exact test), indicating that most H1 hyperDMRs are DME targets (*Figure 2E-H*).

Our results demonstrate that H1 hyperDMRs are primarily caused by the inhibition of DME. However, only 3066 out of 11896 (26%) VC DME targets have significantly more CG methylation in the VC of *pVC::H1* than WT (p<0.001, Fisher’s exact test; *Figure 1—source data 2*), indicating that VC-expressed H1 impedes DME at a minority of its genomic targets. These H1-impeded DME targets are heterochromatic, significantly enriched in H3K9me2 compared with H1-independent DME targets (*Figure 2I*). To further examine the link with heterochromatin, we aligned all VC DME target loci at the most hypomethylated cytosine, and separated them into five groups by H3K9me2 levels (*Figure 2J*). *pVC::H1*-induced hypermethylation peaks where DME-mediated hypomethylation peaks, but is apparent only in the most heterochromatic group (highest H3K9me2) of DME target loci (*Figure 2J*). Taken together, our results demonstrate that developmental removal of H1 from the VC allows DME to access heterochromatin.

### H1 represses transposons via methylation-dependent and independent mechanisms

Given the importance of H1 removal for DME-directed DNA demethylation, we investigated the contribution of H1 to TE activation in the VC. RNA-seq was performed using pollen from the *pVC::H1* line (#2), which showed strong H1 expression in VC (*Figures 2B* and *3A*). 47 out of 114 (41%) VC-activated TEs show significant differential expression (fold change > 2; p<0.05, likelihood ratio test) due to H1 expression in VC (*Figure 3B*). Among these differentially expressed TEs, the overwhelming majority (46; 98%) are repressed (*Figure 3B,C*, *Figure 1—source data 1*). In contrast to the effect of H1 on TE transcription, a much smaller fraction of genes (3%; 89 out of 2845 pollen-expressed genes) is differentially expressed (fold change > 2; p<0.05, likelihood ratio test) between *pVC::H1* and WT (*Figure 3D*). These data indicate that ectopic expression of H1 in the VC preferentially represses TEs.

**Figure 3.**
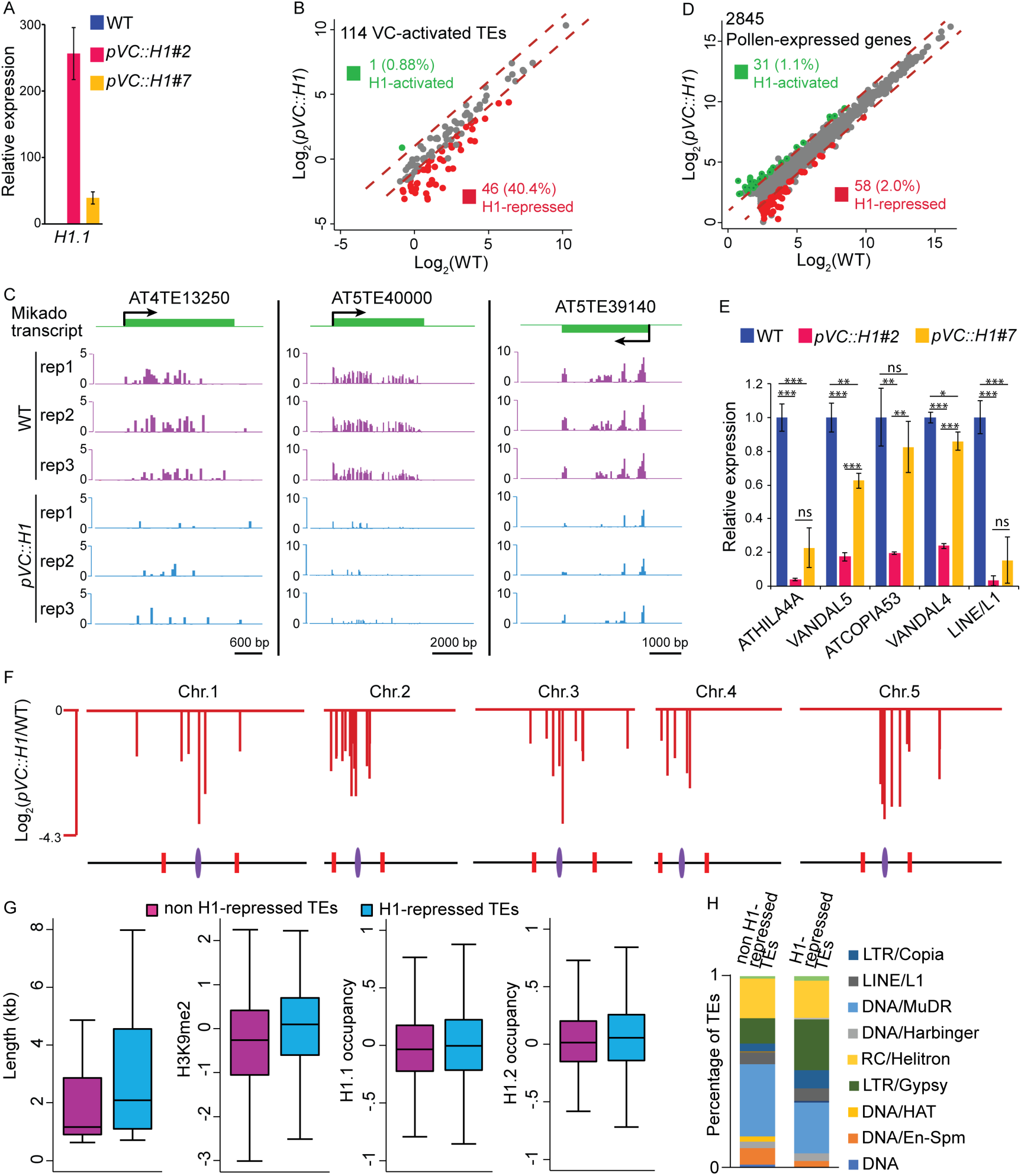
Vegetative-cell-expressed H1 represses heterochromatic TEs in a dosage-dependent manner. *pVC::H1* refers to line #2 except as specified in **A** and **E**. (**A**,**E**) quantitative RT-PCR demonstrating *H1.1* (**A**) or TE (**E**) expression in pollen from WT and two independent *pVC::H1* transgenic lines. Relative expression is calculated by normalizing to WT (WT = 1). Student’s *t* test *p<0.05, **p<0.01, ***p<0.001; ns, not significant; n=3; mean ± SD are shown. (**B**,**D**) Scatter plot illustrating the expression (Log_2_TPM) of TEs or genes in WT and *pVC::H1* pollen. Red and green dots indicate significant down‐ and up-regulation in *pVC::H1* compared to WT (|fold change| > 2, marked by dashed lines; likelihood ratio test p < 0.05), respectively. (**C**) Snapshots showing the expression (Log_2_RPKM) of 3 example H1-repressed TEs in WT and *pVC::H1* pollen. Rep, biological replicate. (**F**) Chromosomal view of H1-repressed TEs, similar to *Figure 1A*. (**G**) Box plots illustrating the length, H3K9me2 enrichment, and H1 occupancy at two groups of VC-activated TEs. Difference between the two datasets compared for each feature is significant (Kolmogorov-Smirnov test *p* < 0.05 for length, and < 0.001 for others). (**H**) Percentages of TEs classified by superfamily.

Quantitative RT-PCR validated our RNA-seq results and confirmed the strong suppression of TEs in *pVC::H1* (*Figure 3E*). Taking advantage of a *pVC::H1* line #7 with weaker H1 expression in pollen (*Figure 3A*), we found H1 represses TE expression in a dosage-dependent manner; as TEs are suppressed to a lesser extent in line #7 compared to the strong line #2 (*Figure 3E*). H1-repressed TEs in the VC are predominantly localized to pericentromeric regions and are overrepresented for LTR retrotransposons, including Gypsy and Copia elements (*Figure 3F-H*). Compared to other VC-activated TEs, the H1-repressed TEs are significantly longer and enriched for H3K9me2 and H1 in somatic tissues (*Figure 3G*), consistent with the observation that H1 precludes DME access to heterochromatin.

In support of the hypothesis that H1 represses VC TE expression by blocking DME, 18 of 46 H1-repressed TEs show significant increase of DNA methylation in at least one sequence context within 300 bp of the TSS in *pVC::H1* (p<0.001, Fisher’s exact test; *Figure 4A,B*). Six more TEs overlap a DME target, which is hypermethylated in *pVC::H1*, within 1 kb of the TSS, and hence may also be suppressed by DME inhibition. However, 22 TEs do not overlap any H1 hyperDMRs within 1 kb of the TSS (*Figure 4A*, marked by asterisks in the lower panel), indicating that their suppression by H1 is not mediated by DNA methylation. Of these, 16 TEs overlap DME targets within 1 kb of TSS. DME maintains access to these TEs in the presence of H1, suggesting their VC demethylation does not rely on the depletion of H1 and their repression in *pVC::H1* is DME-independent; exemplified by AT3TE60310 (*Figure 4C*). Our results demonstrate that H1 overexpression in the VC represses heterochromatic TEs via both DNA methylation-dependent and independent mechanisms.

**Figure 4.**
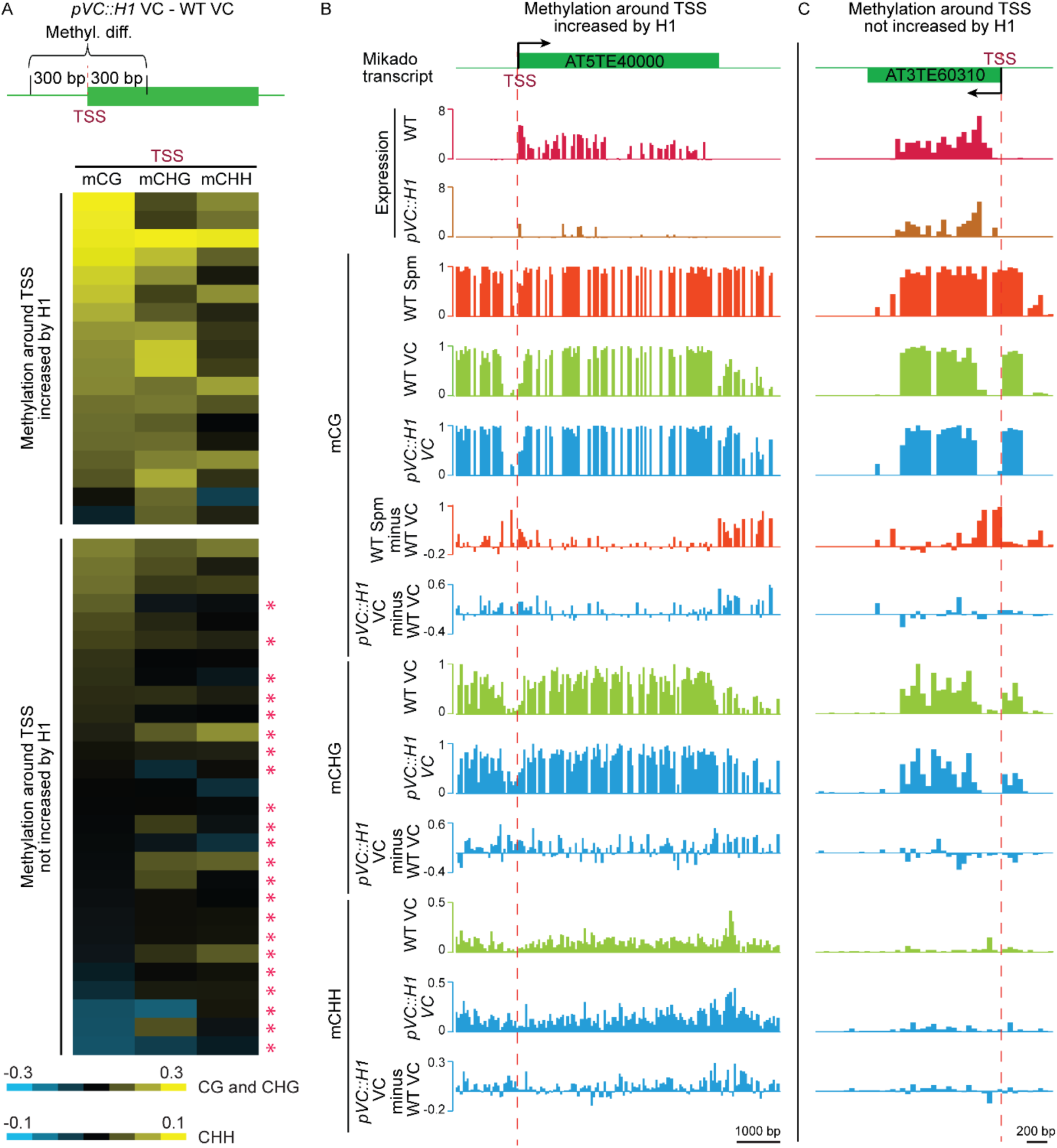
H1 suppresses TEs in the vegetative cell via two mechanisms. (**A**) Heat map demonstrating DNA methylation differences between *pVC::H1* and WT VCs within 300 bp of the TSS of H1-repressed TEs. Asterisks mark TEs whose suppression is not caused by changes in DNA methylation. Data are sorted in descending order based on CG methylation difference for upper and lower panels, respectively. (**B**,**C**) Snapshots showing the expression and DNA methylation of representative TEs suppressed by *pVC::H1* via methylation-dependent (**B**) and ‐independent (**C**) mechanisms, respectively. Spm, sperm.

### Depletion of H1 decondenses heterochromatin during male gametogenesis

H1 depletion and TE activation in the VC are accompanied by loss of cytologically detectable heterochromatin (Baroux et al., 2011; Ingouff et al., 2010; Schoft et al., 2009). We therefore tested whether H1 contributes to heterochromatin condensation in plant cells. Immunostaining of leaf nuclei showed that H1 co-localizes with H3K9me2 in highly-compacted heterochromatic foci, known as chromocenters (*Figure 5A*). Furthermore, we found that chromocenters become dispersed in the nuclei of *h1* mutant rosette leaves (*Figure 5B*). These observations demonstrate that H1 is required for heterochromatin condensation in plants.

**Figure 5.**
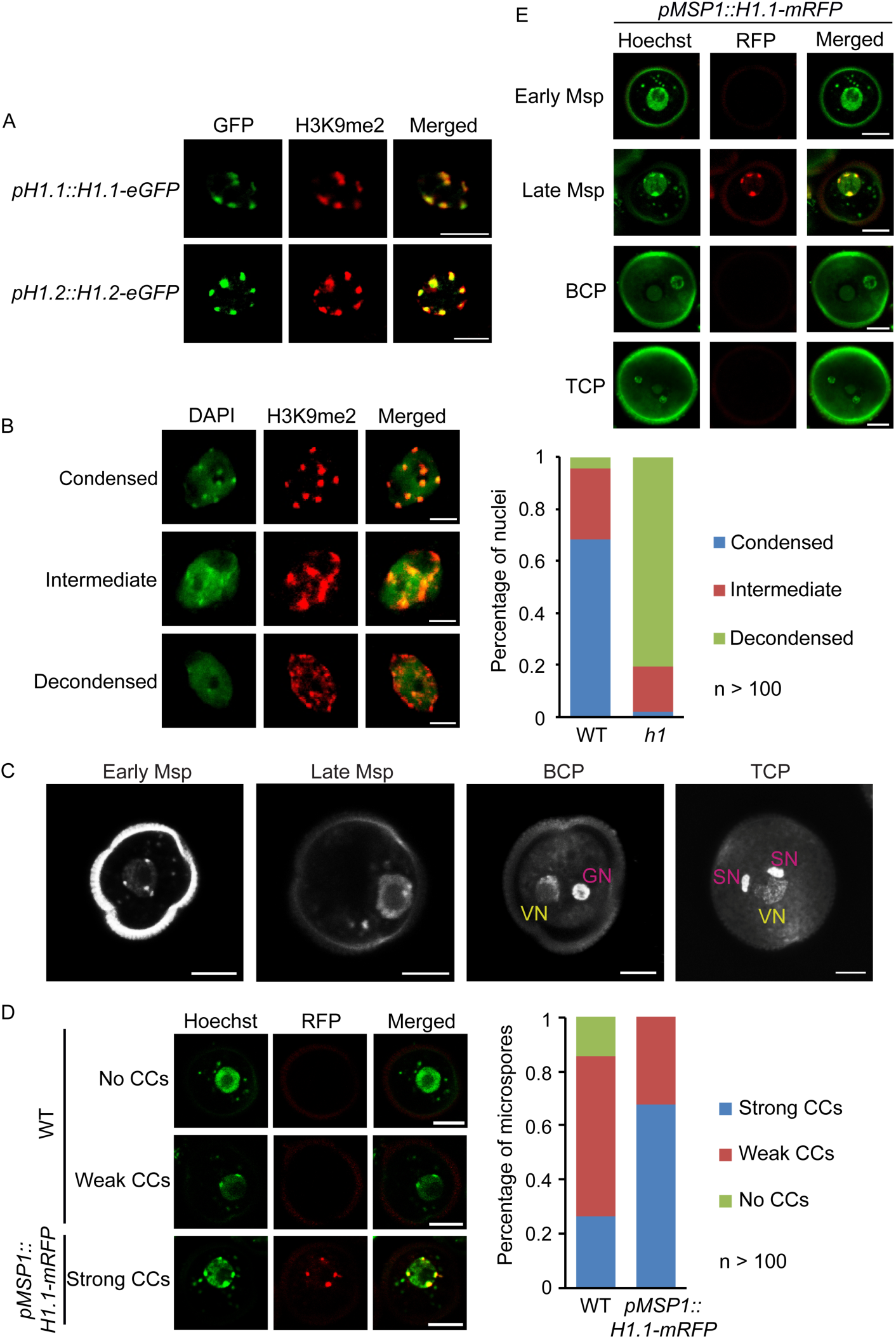
Depletion of H1 decondenses heterochromatin in leaves and late microspores. (**A**) Immunostaining with GFP and H3K9me2 antibodies showing the co-localization of H1 and H3K9me2-enriched chromocenters. (**B**) Examples of leaf nuclei with condensed, intermediate or decondensed chromocenters, and their percentages in WT or the *h1* mutant. Msp, microspore; BCP, bicellular pollen; TCP, tricellular pollen. (**C**) Gradual decondensation of heterochromatin during male gametogenesis in *Arabidopsis*. Micrographs of Hoechst-stained microspores and pollen demonstrate a gradual dispersion of chromocenters in late microspores and subsequently the vegetative nucleus (VN) in pollen. Chromocenters are not detected in the VN of BCP and TCP. GN, generative nucleus. (**D**) Percentages of late microspores with no, weak or strong chromocenters (CCs; examples on the left) in WT and *pMSP1::H1.1-mRFP*, in which H1 colocalizes with the strong CCs. (**E**) H1 is induced only in late microspores in *pMSP1::H1.1-mRFP*, and co-localizes with strong CCs. All bars, 5 μm.

We then examined whether ectopic H1 expression can condense the heterochromatin in VC nuclei. Consistent with previous observations (Baroux et al., 2011; Ingouff et al., 2010; Schoft et al., 2009), no condensed chromocenters were detected in WT VC (*Figure 5C*). *pVC::H1* VC also showed no obvious chromocenters (*Figure 2B*). This suggests either that H1 expression is not strong enough in *pVC::H1*, or other factors are involved in heterochromatin decondensation in the VC.

Heterochromatin decondensation during male gametogenesis seems to be gradual: chromocenters are observed at early microspore stage, but become dispersed in late microspore stage, when H1 is depleted (*Figures 2A* and *5C*). We observed strong and weak chromocenters in 27% and 59%, respectively, of late microspore nuclei, whereas no chromocenters were observed in the VC at either bicellular or tricellular pollen stage (*Figure 5C,D*). The further decondensation of VC heterochromatin after H1 depletion during the late microspore stage suggests the involvement of other factors in the VC. To test whether H1 is sufficient to induce chromatin condensation in microspores, we used the late-microspore-specific *MSP1* promoter (Honys et al., 2006) to drive H1 expression (*pMSP1::H1.1-mRFP*, short as *pMSP1::H1*). In *pMSP1::H1*, we observed strong chromocenters in the majority (68%) of late microspores (*Figure 5D*). H1 expression in *pMSP1::H1* is specific to late microspores, and co-localizes with induced chromocenters (*Figure 5E*). These results show that H1 is sufficient to promote heterochromatic foci in late microspores, thus demonstrating the causal relationship between H1 depletion and the decondensation of heterochromatin.

## Discussion

Epigenetic reactivation of TEs in the VC of flowering plants is an intriguing phenomenon, which is important not only for understanding sexual reproduction, but also for elucidating epigenetic silencing mechanisms. Here we show that *Arabidopsis* VC-activated TEs are heterochromatic, and mostly subject to DME-directed demethylation at their TSS (*Figure 1F*). Given the well-demonstrated role of DNA methylation at the TSS for transcriptional suppression (Barau et al., 2016; Eichten et al., 2012; Hollister & Gaut, 2009; Manakov et al., 2015; Meng et al., 2016), our data demonstrate that DME-mediated demethylation in the VC is the primary cause of TE activation. As DNA demethylation of TEs during reproduction also occurs in rice and maize (Park et al., 2016; Rodrigues et al., 2013; M. Zhang et al., 2014), species that diverged from *Arabidopsis* more than 150 million years ago (Chaw, Chang, Chen, & Li, 2004), our results suggest that TE activation in the VC is prevalent among flowering plants.

DME demethylates about ten thousand loci in the VC and central cell, respectively, however, only half of these loci overlap (Ibarra et al., 2012). It was unclear why DME targets differ in these cell types, but differences in chromatin configuration have been postulated to contribute (Feng et al., 2013). Our finding that the access of DME to heterochromatic TEs in the VC is permitted by the lack of H1 supports this idea. H1 is presumably present in the central cell (Frost et al., 2018) but is absent in the VC (Hsieh et al., 2016), thus rendering heterochromatic TEs more accessible in the VC. Differential distribution of other factors in the VC and central cell, such as histone variant H3.1 (Borg & Berger, 2015; Ingouff et al., 2010), may also affect DME targeting. Consistently, FACT is required for DME activity in the central cell at many loci even in the absence of H1, whereas DME is entirely independent of FACT in the VC (Frost et al., 2018), suggesting the presence of impeding factor(s) other than H1 in the central cell. With distinct chromatin architectures, the vegetative and central cells are excellent systems for understanding how chromatin regulates DNA demethylation.

Our finding that histone H1 affects DME activity adds to the emerging picture of H1 as an important and complex regulator of eukaryotic DNA methylation. H1 depletion causes local hypomethylation in mouse cells (Fan et al., 2005) and extensive hypermethylation in the fungi *Ascobolus immersus* (Barra, Rhounim, Rossignol, & Faugeron, 2000) and *Neurospora crassa* (Seymour et al., 2016). In *Arabidopsis*, loss of H1 causes global heterochromatic hypermethylation in all sequence contexts by allowing greater access of DNA methyltransferases (Lyons & Zilberman, 2017; Zemach et al., 2013). Our results suggest that H1 may also influence DME-homologous demethylases that control methylation in somatic tissues (He et al., 2011). By regulating both methylation and demethylation, H1 may serve as an integrator of methylation pathways that tunes methylation up or down depending on the locus.

Our data also indicate that the regulatory functions of H1 extend beyond DNA methylation in plants. Activated TEs in the VC can be categorized into four groups, based on the mechanism of their activation (*Figure 6*). TEs in Group I are the least heterochromatic and their activation is dependent on DME but not H1 (*Figures 3D* and *6*). Group II comprises TEs in which H1 absence is required for DME demethylation and activation (*Figure 6*). For TEs in Group III, H1 depletion and DME demethylation are both required for activation, but DME activity is not affected by H1 (*Figure 6*). Group IV TEs are activated by H1 depletion and are not targeted by DME (*Figure 6*). Groups III and IV demonstrate that H1 can silence TEs independently of DNA methylation. Group III also demonstrates that DNA methylation and H1 cooperate to suppress TE expression in plants. Thus, H1 regulates TEs via DNA methylation-dependent and-independent mechanisms.

**Figure 6.**
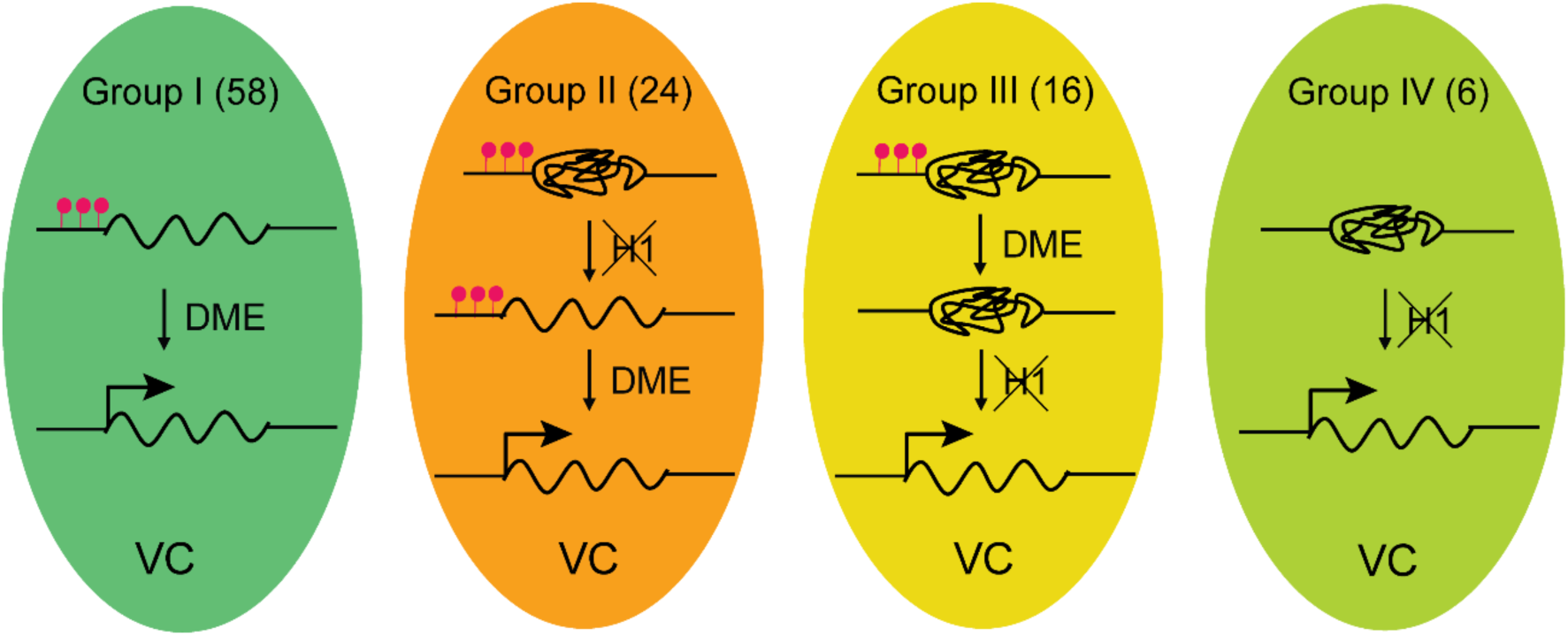
Model depicting four mechanisms underlying TE activation in the VC. The number of TEs in each group is shown on the top. Significantly less heterochromatic than TEs in other groups (*Figure 3G*), Group I TEs are activated by DME-directed DNA demethylation. Group II TEs rely on H1 depletion to allow DME demethylation and activation. Group III TEs are demethylated by DME but require H1 depletion to allow transcription (ie. *pVC::H1* represses these TEs without affecting DME). Group IV TEs are not demethylated by DME; their activation is solely dependent on the depletion of H1. TEs belong to each group are listed in *Figure 1—source data 1*. Red lollipops denote DNA methylation.

During the ongoing arms race between TEs and their hosts, it may be difficult to determine whether TE expression represents temporary TE triumphs or is domesticated by the host to serve a function. TE activation in the VC – a cell that engulfs the male plant gametes – has been proposed as a defense strategy, which generates small RNAs that enhance TE silencing in sperm (Calarco et al., 2012; Ibarra et al., 2012; Martinez et al., 2016; Slotkin et al., 2009). However, TEs can also use companion cells as staging grounds for invasion of the gametes (Wang, Dou, Moon, Tan, & Zhang, 2018). Our demonstration that programmed DME demethylation, which is facilitated by developmental heterochromatin decondensation, is the predominant cause of VC TE activation is consistent with a defensive, host-beneficial model. Nonetheless, the alternative TE-driven model is also plausible. DME demethylation regulates genes and is important for pollen fertility (Choi et al., 2002; Ibarra et al., 2012; Schoft et al., 2011). Our data show that developmental H1 depletion is also important for pollen fertility. Therefore, at least some TEs may be hijacking an essential epigenetic reprogramming process. TE activation in the VC may facilitate both host defense and transposition, with the balance specific to each TE family and changing over evolutionary time. The effects of VC TE activation on TE proliferation in the progeny may warrant investigation, particularly in out-crossing species with aggressive TEs and in natural populations.

## Materials and Methods

### Plant materials and growth conditions

*A. thaliana* plants were grown under 16h light/ 8h dark in a growth chamber (20°C, 80% humidity). All plants used are of the Col-0 ecotype. *pH1.1::H1.1-eGFP*, *pH1.2::H1.2-eGFP* and the *h1* (*h1.1 h1.2* double) mutant lines were described previously (She et al., 2013; Zemach et al., 2013). *pLAT52::H1.1-mRFP* and *pMSP1::H1.1-mRFP* were constructed with MultiSite Gateway System into the destination vector pK7m34GW (Invitrogen). The BP clones pDONR-P4-P1R-*pLAT52* and pDONR-P2R-P3-mRFP were kindly provided by Prof. David Twell (Leicester University, UK) (Eady et al., 1994). *MSP1* promoter was cloned into pDONR-P4-P1R as described previously (Honys et al., 2006). WT plants were transformed via floral dip (Clough & Bent, 1998), and T2 or T3 plants homozygous for the transgene were used in this study.

### Pollen extraction, RNA sequencing and quantitative RT-PCR

Open flowers were collected for pollen isolation in Galbraith buffer (45 mM MgCl2, 30 mM sodium citrate, 20 mM MOPS, 1% Triton-X-100, pH7.0) by vortexing at 2000 rpm for 3 min. The crude fraction was filtered through a 40 μm cell strainer to remove flower parts, and subsequently centrifuged at 2600 g for 5 min to obtain pollen grains. RNA was extracted from pollen grains with RNeasy Micro Kit (Qiagen) following manufacturer’s instructions. RNA-sequencing libraries were prepared using Ovation RNA-seq Systems 1-16 for Model Organisms (Nugen Technologies), and sequenced on the Hiseq 2500 (Illumina) instrument at the UC Berkeley Vincent J. Coates Genomics Sequencing Laboratory. Quantitative RT-PCR (qRT-PCR) was performed as described previously (Walker et al., 2018), and *TUA2* was used as an internal control. Primers for qRT-PCR are listed in Supplementary file 2.

**Supplementary file 2.**
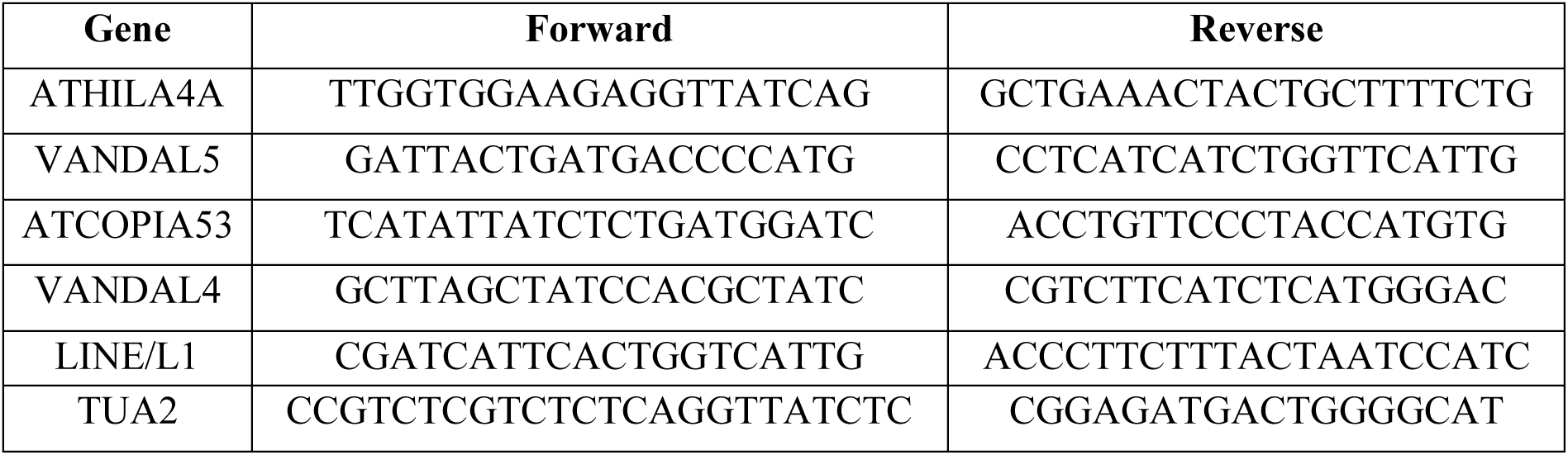
List of primers for quantitative RT-PCR.

### RNA-seq analysis

TE transcript annotation was created using RNA-seq data from four biological replicates of pollen. Tophat2, Hisat, and STAR were used to align RNA-seq reads to the TAIR10 genome, and transcripts were assembled using CLASS2, StringTie, and Cufflinks, respectively. Assembled transcripts were selected by Mikado using default options except that the BLAST and Transdecoder steps were disabled (Venturini et al., 2018). As a result, 21381 transcripts (called superloci; GSE120519) were identified.

To identify VC-activated TEs, we first refined the list of superloci by selecting those overlapping with TAIR10 TE annotation. Subsequently to eliminate TE-like genes from the refined list, superloci with CG methylation less than 0.7 in rosette leaves (Stroud et al., 2014; Stroud, Greenberg, Feng, Bernatavichute, & Jacobsen, 2013) were excluded. This gave rise to an annotation of pollen TE transcripts, which was combined with TAIR10 gene annotation for Kallisto analysis (Bray, Pimentel, Melsted, & Pachter, 2016). RNA-seq data from pollen (this study) and rosette leaves (Walker et al., 2018), each including three biological replicates, were processed using Kallisto and Sleuth (Bray et al., 2016; Pimentel, Bray, Puente, Melsted, & Pachter, 2017). TEs that are transcribed at least 5 times more in pollen than leaves (with p < 0.05, likelihood ratio test) are considered as activated in the VC (refer to *Figure 1—source data 1* for the list of VC-activated TEs). A total of 2845 genes were found to be expressed in pollen with TPM (transcripts per million) more than 5 in the Kallisto output (data used in *Figure 3D*).

To identify TEs and genes that are suppressed by H1 in the VC, we analyzed RNA-seq data from WT and *pLAT52::H1.1-mRFP line #2* (short as *pVC::H1* unless specified otherwise) pollen using Kallisto and Sleuth as described above. Significant differential expression was defined with a fold change at least 2 and a p-value less than 0.05. H1-repressed TEs were listed in *Figure 1—source data 1*.

### Whole-genome bisulfite sequencing and analysis

Vegetative and sperm nuclei were isolated via FACS as described previously (Ibarra et al., 2012). Bisulfite-sequencing libraries were prepared as previously described (Walker et al., 2018). Sequencing was performed on Hiseq 2500 (Illumina) at the UC Berkeley Vincent J. Coates Genomics Sequencing Laboratory, Hiseq 4000 (Illumina) at Novogene Ltd. and Harvard University, and Nextseq 500 (Illumina) at Cambridge University Biochemistry Department and the John Innes Centre. Sequenced reads (100, 75, or 50 nt single-end) were mapped to the TAIR10 reference genome, and cytosine methylation analysis was performed as previously described (Ibarra et al., 2012).

### Identification of DME targets and H1 hyperDMRs in the VC

As all CG hypomethylation in the VC in comparison to sperm is caused by DME (Ibarra et al., 2012), we identified VC DME targets via detecting CG differentially methylated regions (DMRs) that are hypermethylated in sperm in comparison to the VC. DMRs were identified first by using MethPipe (settings: p = 0.05 and bin = 100) (Song et al., 2013), and subsequently retained if the fractional CG methylation across the whole DMR was at least 0.2 higher in sperm than the VC. The refined DMRs were merged to generate larger DMRs if they occurred within 300 bp. Finally, merged DMRs were retained if they cover at least 100 bp, and the fractional CG methylation across the whole DMR was significantly (Fisher’s exact test p < 0.01) and substantially (>0.2) higher in sperm than the VC. This resulted in the identification of 11896 VC DME targets (*Figure 1—source data 2*).

H1 hyperDMRs were identified using the same criteria, except comparing CG methylation in VCs from *pVC::H1* and WT. In total, 2964 H1 hyperDMRs were identified (*Figure 2—source data 1*).

### Box plots

Box plots compare the enrichment of genomic or chromatin features among TEs (*Figure 1B*, *3G*, *Figure 1—figure supplement 1B*) or VC DME targets (*Figure 2I*) as described in corresponding figure legends. ChIP-seq data for H3K9me2 (Stroud et al., 2014), and ChIP-chip data for H1 (Rutowicz et al., 2015), H3K27me3 (Kim, Lee, Eshed-Williams, Zilberman, & Sung, 2012), and other histone modifications (Roudier et al., 2011) were used.

### Density plots

All DNA methylation kernel density plots compare fractional methylation within 50-bp windows. We used windows with at least 20 informative sequenced cytosines and fractional methylation of at least 0.5 (*Figure 2D*, *Figure 2—figure supplement 2*) or 0.7 (*Figure 2E*) for CG context, and 0.4 and 0.1 for CHG and CHH context, respectively, in at least one of the samples being compared.

### Meta analysis (ends analysis)

Ends analysis for TEs and genes was performed as described previously (Ibarra et al., 2012). Similarly, ends analysis of TE transcripts was performed using the annotation of VC-activated TEs described above (*Figure 1—source data 1*). DNA methylation data from (Ibarra et al., 2012) was used.

In *Figure 2J*, DME sites were aligned at the most demethylated cytosine, and average CG methylation levels for each 10-bp interval at both sides were plotted. To identify individual hypomethylation sites created by DME, we first obtained the 50-bp windows with a CG methylation difference larger than 0.5 between sperm and VC (sperm – VC > 0.5 and Fisher’s exact test p < 0.001). Windows were then merged if they occurred within 200 bp. Merged windows were retained for further analysis if the fractional CG methylation across the whole site was 0.2 greater in sperm than VC (sperm – VC > 0.2 and Fisher’s exact test p < 0.0001). This resulted in 13610 DME sites, which were separated into five groups according to H3K9me2 level (Stroud et al., 2014): < 2.5, 2.5-4.3, 4.3-6.5, 6.5-10.5, and > 10.5 (*Figure 2J*). The most demethylated cytosine within each site was identified if it had the greatest differential methylation in sperm than VC among cytosines in the CG context (sperm – VC > 0.2, and Fisher’s exact test p < 0.001) and was sequenced at least 10 times.

### DNA methylation analysis of H1-repressed TEs

Differential methylation at a 600-bp region centered upon the TSS of H1-repressed TEs was calculated between VCs of *pVC::H1* and WT (*Figure 4A*). TEs whose differential methylation is significant (Fisher’s exact test p < 0.001) and larger than 0.2 (in CG context), 0.1 (in CHG context), or 0.05 (in CHH context) are illustrated in the upper panel in *Figure 4A*.

### Confocal and scanning electron microscopy

Microspores and pollen were isolated as described previously (Borges et al., 2012), stained with Hoechst or DAPI, and examined under a Leica SP8 confocal microscope. Scanning electron microscopy was performed on a Zeiss Supra 55 VP FEG.

### Immunofluorescence

Immunofluorescence was performed as described previously with small modifications (Yelagandula et al., 2014). Rosette leaves from 3-week-old plants were fixed in TRIS buffer with 4% paraformaldehyde (10 mM Tris-HCl pH 7.5, 10 mM EDTA, 100 mM NaCl) for 20 min. After being washed with TRIS buffer twice, the fixed leaves were chopped with razor blades in 1 mL of lysis buffer (15 mM Tris pH 7.5, 2 mM EDTA, 0.5 mM spermine, 80 mM KCl, 20 mM NaCl, 0.1% Triton X-100) and filtered through a 35 μm cell strainer. Nuclei were pelleted via centrifugation at 500 g for 3 min and resuspended in 100 μL of lysis buffer. Next, 10 μL was spotted onto coverslips, air-dried, and post-fixed in PBS with 4% paraformaldehyde for 30 min. After being washed with PBS twice, coverslips were incubated in blocking buffer (PBS with 1% BSA) at 37°C for 30 min and then incubated in blocking buffer with primary antibodies at 4°C overnight (Mouse anti-H3K9me2 Abcam ab1220, 1:100; Rabbit anti-GFP Abcam ab290, 1:100). After being washed with PBS three times, coverslips were incubated in PBS with secondary antibodies at 37°C for 30 min, and then washed with PBS three times again before being counterstained and mounted in Vectashield mounting media with DAPI (Vector H-1200).

## Acknowledgements

We thank David Twell for the pDONR-P4-P1R-pLAT52 and pDONR-P2R-P3-mRFP vectors, the John Innes Centre Bioimaging Facility (Elaine Barclay and Grant Calder) for their assistance with microscopy, and the Norwich BioScience Institute Partnership Computing infrastructure for Science Group for High Performance Computing resources. This work was funded by a Biotechnology and Biological Sciences Research Council (BBSRC) David Phillips Fellowship (BBL0250431) and Grant to Exceptional Researchers by the Gatsby Charitable Foundation to X.F.

## Author contributions

S.H. and X.F. designed the study and wrote the manuscript, S.H. and J.Z. performed the experiments, and S.H. and M.V. analyzed the data.

## Author information

Sequencing data have been deposited in GEO (GSE120519). The authors declare no competing financial interests. Correspondence and requests for materials should be addressed to X.F.

---

**Figure 1—figure supplement 1.**
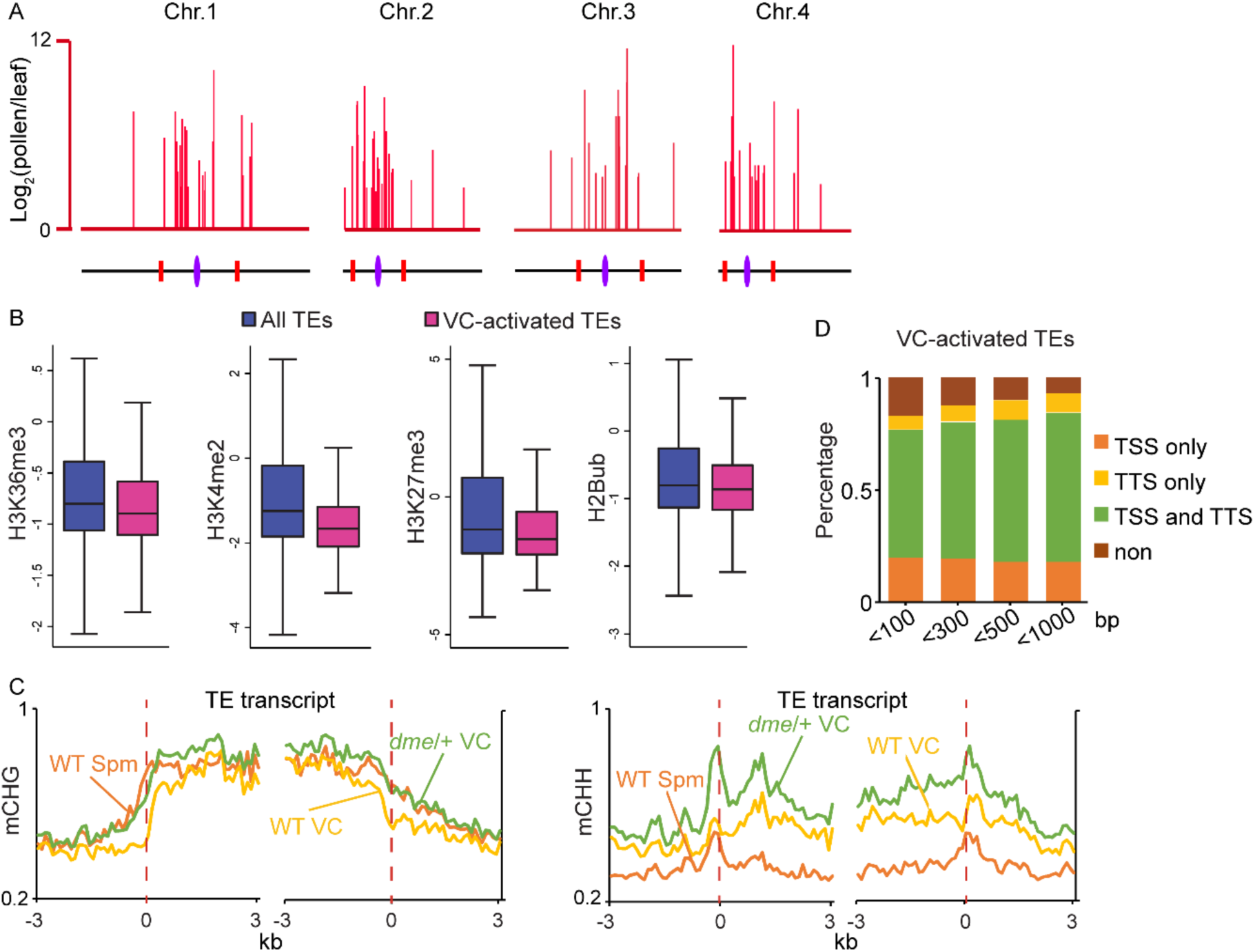
VC-activated TEs are heterochromatic and demethylated by DME. (**A**) Chromosomal view of VC-activated TEs, similar to *Figure 1A*. (**B**) Box plots showing the enrichment of euchromatic histone modifications at TEs, similar to *Figure 1B*. Difference between the two datasets compared for each feature is significant (Kolmogorov-Smirnov test *p* < 0.001). H2Bub, H2B ubiquitination. (**C**) Ends analysis of VC-activated TEs, similar to *Figure 1D*. (**D**) Percentages of VC-activated TEs with TSS and/or TTS within indicated distances of VC DME targets.

**Figure 2—figure supplement 1.**
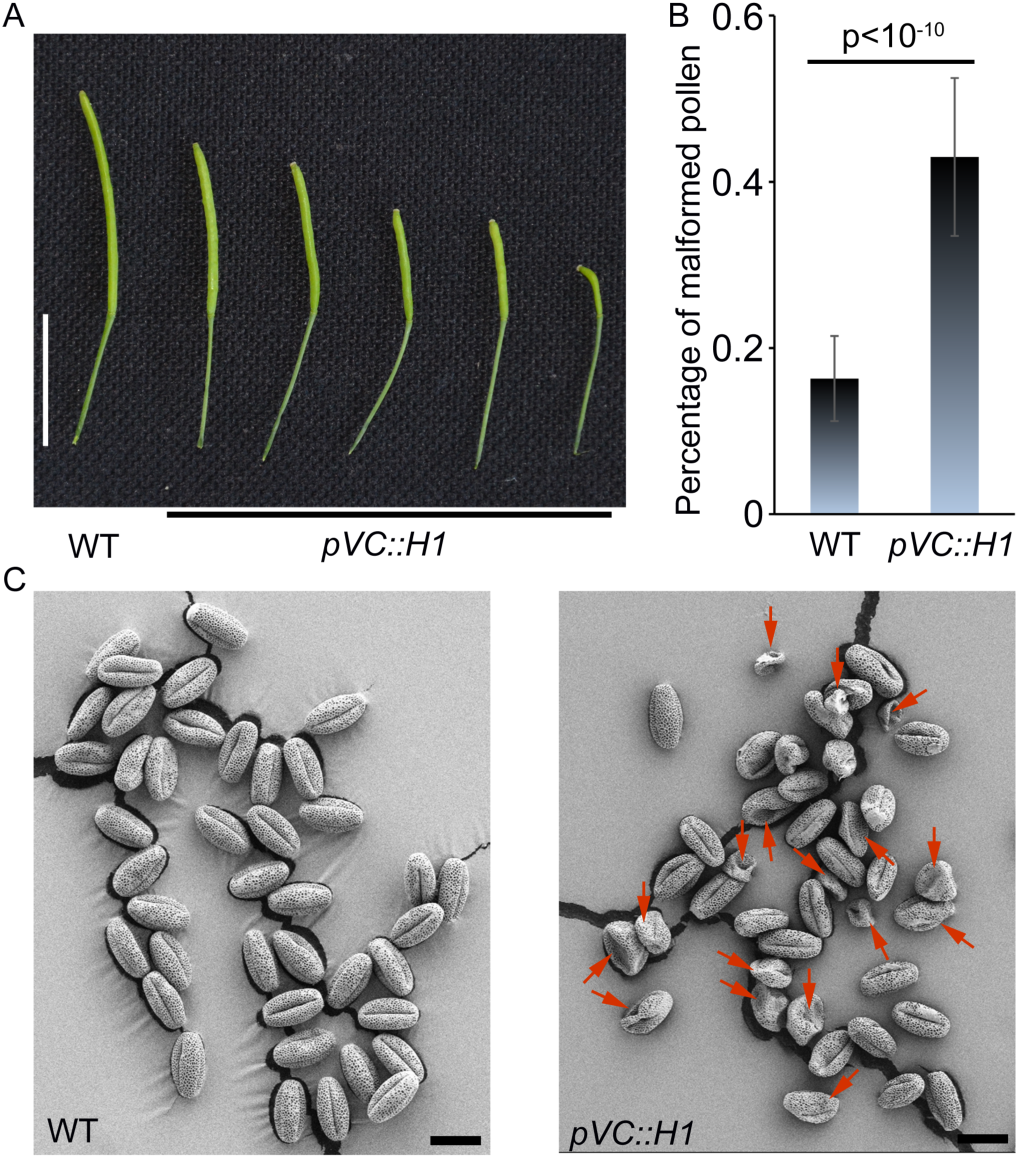
H1 ectopic expression in the vegetative cell causes pollen defect and reduced fertility. *pVC::H1* (*pLAT52::H1-mRFP* line #2) plants show reduced silique length (**A**) and an increased proportion of malformed pollen grains (**B**), which are indicated by red arrows in the SEM image (**C**). Student’s *t* test p<10^−10^; n=17; mean ± SD are shown. Bar (**A**), 1 cm; Bar (**C**), 20 μm.

**Figure 2—figure supplement 2.**
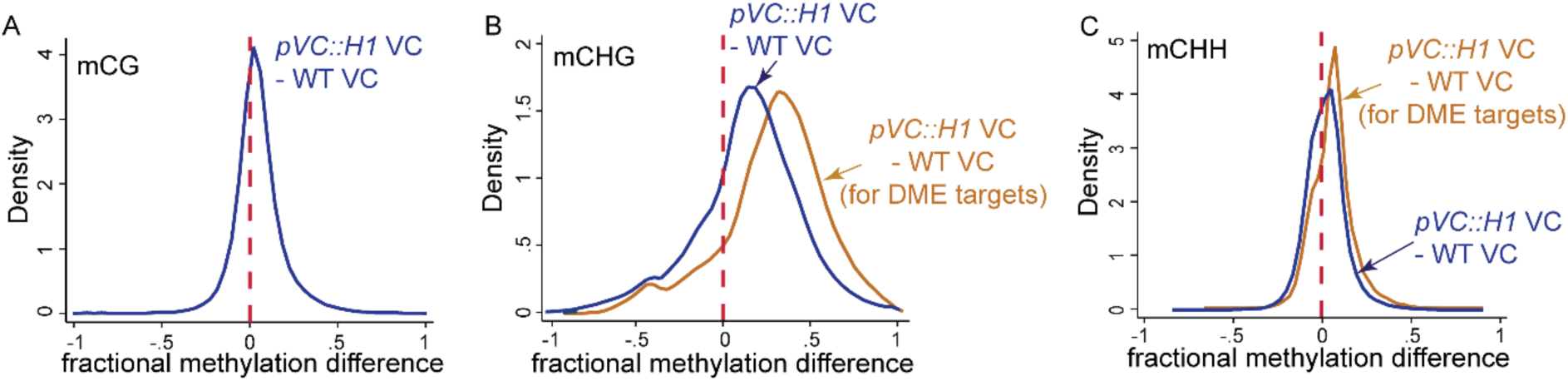
H1 ectopic expression in the vegetative cell causes DNA hypermethylation at DME targets. Kernel density plots showing frequency distribution of methylation differences between VCs from *pVC::H1* and WT in all 50-bp windows (blue traces) and windows overlapping VC DME targets (orange traces).

